# A rate threshold mechanism regulates MAPK stress signaling and survival

**DOI:** 10.1101/155267

**Authors:** Amanda N. Johnson, Guoliang Li, Hossein Jashnsaz, Alexander Thiemicke, Benjamin K. Kesler, Dustin C. Rogers, Gregor Neuert

## Abstract

Cells are exposed to changes in extracellular stimulus concentration that vary as a function of rate. However, the effect of stimulation rate on cell behavior and signaling remains poorly understood. Here, we examined how varying the rate of stress application alters budding yeast cell viability and mitogen-activated protein kinase (MAPK) signaling at the single-cell level. We show that cell survival and signaling depend on a rate threshold that operates in conjunction with a concentration threshold to determine the timing of MAPK signaling during rate-varying stimulus treatments. We also discovered that the stimulation rate threshold is sensitive to changes in the expression levels of the Ptp2 phosphatase, but not of another phosphatase that similarly regulates osmostress signaling during switch-like treatments. Our results demonstrate that stimulation rate is a regulated determinant of signaling output and provide a paradigm to guide the dissection of major stimulation rate-dependent mechanisms in other systems.

## Main Text

All cells employ signal transduction pathways to respond to physiologically relevant changes in extracellular stressors, nutrient levels, hormones, morphogens, and other stimuli that vary as functions of both concentration and rate in healthy and diseased states^1–7^. Switch-like “instantaneous” changes in the concentrations of stimuli in the extracellular environment have been widely used to show that the strength of signaling and overall cellular response are dependent on the stimulus concentration, which in many cases needs to exceed a certain threshold^8, 9^. Although studies have shown that the rate of stimulation can also influence signaling output in a variety of pathways^10–17^, it is not clear how cells integrate information conveyed by changes in both the stimulation rate and concentration in determining whether or not to commence signaling.

Recent investigations have demonstrated that stimulation rate can be a determining factor in signal transduction. In contrast to switch-like perturbations, which trigger a broad set of stress response pathways, slow stimulation rates activate a specific response to the stress applied in *Bacillus subtilis* cells^10^. Meanwhile, shallow morphogen gradient stimulation fails to activate developmental pathways in mouse myoblast cells in culture, even when concentrations sufficient for activation during pulsed treatment are delivered^12^. These observations raise the possibility that stimulation profiles must exceed a set minimum rate or rate threshold to achieve signaling activation. Although such rate thresholds would help cells decide if and how to respond to dynamic changes in stimulus concentration, the possibility of signaling regulation by a rate threshold has never been directly investigated in any system. Further, no study has experimentally examined how stimulation rate requirements impact cell phenotype or how cells molecularly regulate the stimulation rate required for signaling activation. As such, the biological significance of any existing rate threshold regulation of signaling remains unknown.

The budding yeast *Saccharomyces cerevisiae* high osmolarity glycerol (HOG) pathway provides an ideal model system for addressing these issues (Fig. 1a). The evolutionarily conserved mitogen-activated protein kinase (MAPK) Hog1 serves as the central signaling mediator of this pathway^18–20^. It is well established that instantaneous increases in osmotic stress concentration induce Hog1 phosphorylation, activation, and translocation to the nucleus^19, 21–29^. Activated Hog1 governs the majority of the cellular osmoadaptation response that enables cells to survive^21, 30, 31^. Multiple apparently redundant MAPK phosphatases dephosphorylate and inactivate Hog1, which, along with the termination of upstream signaling after adaptation, results in its return to the cytosol (Fig. 1a)^21, 23, 24, 32–38^. Because of this behavior, time-lapse analysis of Hog1 nuclear enrichment in single cells has proven an excellent way to monitor signaling responses to dynamic stimulation patterns in real-time^25–29, 39, 40^. Further, such assays have been readily combined with traditional growth and molecular genetic approaches to link observed signaling responses with cell behavior and signaling pathway architecture^26–28^.

**Fig. 1.**
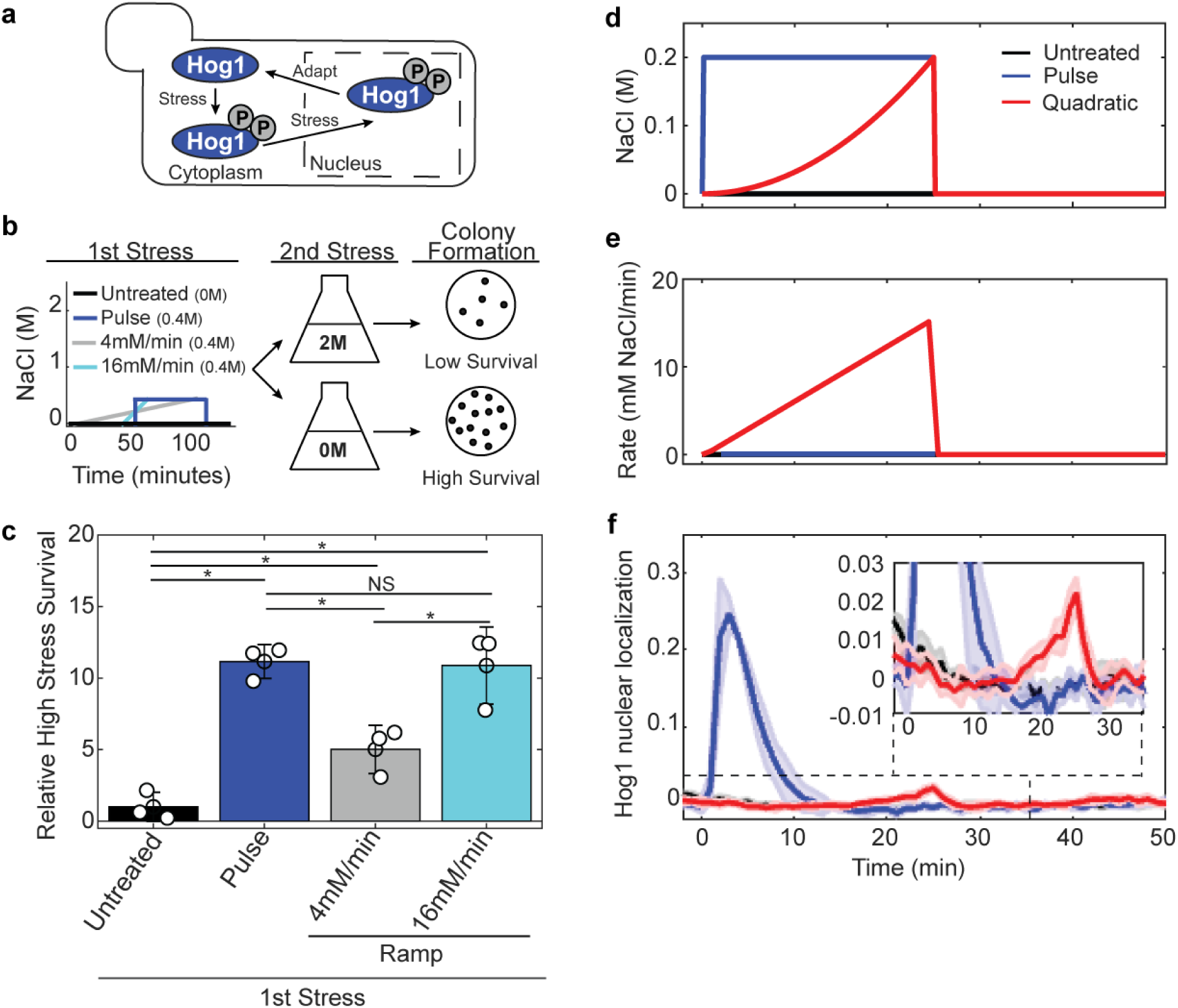
Hog1 signaling and cell survival are sensitive to the rate of preconditioning osmotic stress application. (**a**) Schematic of the budding yeast high osmolarity glycerol (HOG) response. (**b**) Preconditioning protection assay workflow indicating first stress treatments (left plot), high-stress exposure (middle panel), and colony formation readout (right panel). (**c**) High-stress survival as a function of each first treatment relative to the untreated first stress condition. Bars and errors are means and standard deviation from three biological replicates. * = statistically significant by Kolmogorov-Smirnov (KS) test (p < 0.05). NS = not significant. (**d**) Treatment concentration over time. (**e**) Treatment rate over time. (**f**) Hog1 nuclear localization during the treatments depicted in “d” and “e”. Insert highlights localization pattern in the quadratic treated sample. Lines represent means and shaded error represents the standard deviation from 3-4 biological replicates.

Here, we use systematically-designed osmotic stress treatments imposed at varying rates of increase to show that a rate threshold condition regulates yeast high-stress survival and Hog1 MAPK signaling. We demonstrate that only stimulus profiles that satisfy both this rate threshold condition and a concentration threshold condition result in robust signaling. We go on to show that the protein tyrosine phosphatase Ptp2, but not the related Ptp3 phosphatase, serves as a major rate threshold regulator. By expressing *PTP2* under the control of a series of different enhancer-promoter DNA constructs, we demonstrate that changes in the level of Ptp2 expression can alter the stimulation rate required for signaling induction. These findings establish rate thresholds as a critical and regulated component of signaling biology akin to concentration thresholds.

## Results

### Stimulus treatment rate affects yeast high osmotic stress survival and Hog1 nuclear translocation pattern

We hypothesized that a rate threshold governs the budding yeast HOG pathway. The existence of such a threshold would place a minimum stimulation rate on osmoadaptation. Shallow rates of stimulation that fall below this minimum would not result in a robust osmoadaptive response, even if the same stimulus concentration triggered a response when applied at a faster rate.

To test this hypothesis, we first took advantage of the fact that pre-exposure to a mild pulsed stress treatment increases the stress tolerance of yeast through induction of the osmoadaptive response^41–43^. This increased tolerance enables cells to survive a second, otherwise lethal, high-stress pulse treatment^41, 42^. This preconditioning protection paradigm enables us to examine the rate-dependence of mild stress responses independent of whether the stresses themselves differentially impact cell fitness.

Cells were initially exposed to preconditioning osmostresses (400 mM NaCl final concentration, equivalent integrated osmolarity) delivered at different rates. The impact of these preconditioning treatments on high-stress susceptibility was determined by comparing the viability of cells after a second transient high-stress treatment and no stress treatment (2 M and 0 M NaCl) (Fig. 1b). No substantial difference in growth was observed among the preconditioned cultures immediately following the initial treatments (Supplemental Fig. 1). However, the rate of preconditioning stress delivery profoundly influenced cell survival after the second high-stress; cells exposed to a slow rate treatment [4 mM NaCl/min (grey)] were significantly more sensitive to a high second stress than cells exposed to faster rate treatments [16 mM NaCl/min (cyan) or pulse (blue)] (Fig. 1c).

To investigate the mechanism(s) underlying the priming of high second stress survival in response to variations in preconditioning stimulation rate, we tracked the nuclear enrichment of Hog1-yellow fluorescent protein (YFP), a proxy for canonical salt-stress signaling activation, during rate-varying stress treatments via single-cell time-lapse microscopy. Because previously published studies show that Hog1 nuclear translocation follows changing external stress patterns in real-time^28, 29, 39^, we began probing the relationship between stimulation rate and translocation using a stress treatment applied as a quadratic function (linearly increasing rate over time) (Fig. 1d, e). Compared to pulse treatment, quadratic treatment resulted in an unprecedented delayed and diminished Hog1 nuclear accumulation response (Fig. 1f). Stimulation rate thus strongly influences signaling as well as high-stress survival phenotype, consistent with expectations based on rate threshold regulation of the HOG pathway.

### A rate threshold regulates Hog1 nuclear enrichment

The Hog1 quadratic stimulation response pattern could be related to either known stimulus concentration threshold requirements^23^ or stimulation rate, since both are changing over time (Fig. 2a, b). To investigate these relationships, Hog1 nuclear localization during the quadratic stress gradient was reanalyzed separately as a function of either the treatment concentration or rate. The concentration and rate corresponding with nuclear enrichment were extrapolated using two intersecting lines fitting Hog1 nuclear localization baseline and accumulation increase for each independent experiment. The thresholds for Hog1 translocation were 117.5 ± 34.5 mM NaCl and 12.1 ± 1.6 mM NaCl/min, respectively (Fig. 2a, b). Surprisingly, the concentration associated with nuclear accumulation was much higher than the 50-70 mM salt expected based on published switch-like treatment responses^23, 26^, a result that raises the possibility that the rate of stress change drives Hog1 nuclear enrichment pattern during quadratic stimulation.

**Fig. 2.**
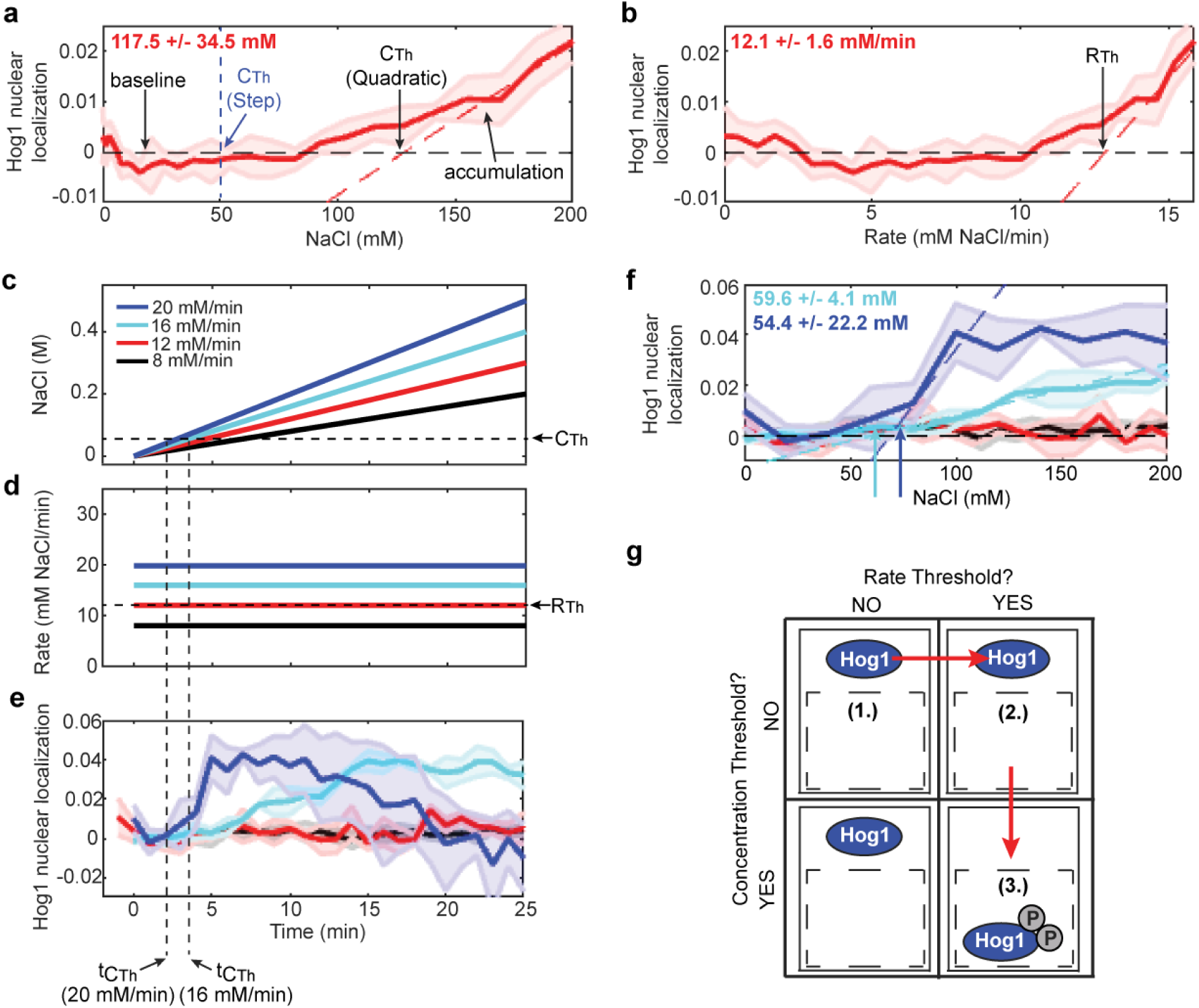
A rate threshold condition regulates signaling nuclear localization. (**a**) Quadratic signaling response from Fig. 1f plotted as a function of treatment concentration. The intersection points between the dashed lines representing Hog1 nuclear localization baseline (black) and accumulation pattern either expected for step treatments (blue) or measured in quadratic treatment (red) indicate concentration thresholds (CTh). The apparent quadratic CTh is listed in bold red font. (**b**) Quadratic treatment response from Fig. 1F plotted as a function of treatment rate. A rate threshold (RTh) (bold red font) was identified via the approach used for CTh determination. (**c**) NaCl concentration change during ramp treatments. CTh is marked. (**d**) NaCl rate during ramp treatments. RTh is indicated. (**e**) Signaling response to ramp treatments plotted as a function of time. (**f**) Signaling response to ramp treatments plotted as a function of treatment concentration. The concentrations corresponding with Hog1 nuclear localization in the ramps where significant activation was detected are indicated in bold font in the top left corner of the plot color-coded to match the corresponding treatment. The concentration corresponding with Hog1 nuclear localization calculated by combining the data from all the biological replicates in both activating ramps was 57 +/-14.5 mM. (**g**) AND logic model indicating the order of condition completion for the ramps exceeding RTh.

To systematically test if ∼12 mM NaCl/min represents an unreported rate threshold requirement for signaling, we exposed cells to linear ramp treatments delivered at rates above, at, and below 12 mM NaCl/min (Fig. 2c, d). Only ramps with rates greater than 12 mM NaCl/min resulted in detectable, albeit slightly delayed, Hog1 translocation (Fig. 2e). This result directly demonstrates that stimulus profiles must exceed a rate threshold to induce Hog1 nuclear enrichment.

### Hog1 translocation requires both the rate and concentration thresholds

If stimulation rate solely determines signaling response, all ramps with gradients steeper than the rate threshold should bring about Hog1 translocation at the onset of treatment. Instead, we observed a delay in translocation in the 16 mM NaCl/min and 20 mM NaCl/min ramp treatments (Fig. 2e). Previously published data also shows a lag in nuclear translocation during ramp treatment^27^, although the corresponding report does not discuss this observation. Reanalyzing translocation during our ramp treatments that resulted in Hog1 nuclear localization as a function of concentration revealed that the timing of translocation in both activating ramps was associated with 57 ± 14.5 mM NaCl (Fig. 2f). This value agrees with the anticipated concentration threshold of 50-70 mM salt.

These results suggest that Hog1 accumulates in the nucleus only when stress stimulations exceed both the rate and concentration thresholds (Fig. 2g). Yeast cell stress signaling responses thus appear to follow AND logic where the output (signaling) occurs only if both input conditions (the rate and concentration thresholds) are met for a given stimulation profile. In this model, neither steps that occur at an infinite rate but fall below the concentration threshold nor slow increases in NaCl that exceed the concentration threshold effectively trigger signaling. On the other hand, for those stimulations where there is a difference in the time each threshold condition is satisfied, Hog1 translocation corresponds with whichever condition was met last. The activating ramps used in these studies exceed the rate threshold before the concentration threshold such that the timing of Hog1 translocation corresponds with the concentration threshold (Fig. 2c-g). In our quadratic stimulation, the order in which these conditions are met is flipped and nuclear accumulation corresponds with the rate threshold instead (Fig. 3a-c, i). The AND logic model thus explains how quadratic stimulations can result in translocation at comparable stimulation rates but different stimulus concentrations (Fig. 3d-i), and how fast rate treatments (e.g. pulses and very fast ramps) which promptly satisfy both threshold conditions induce rapid activation (Fig. 4a-g).

**Fig. 3.**
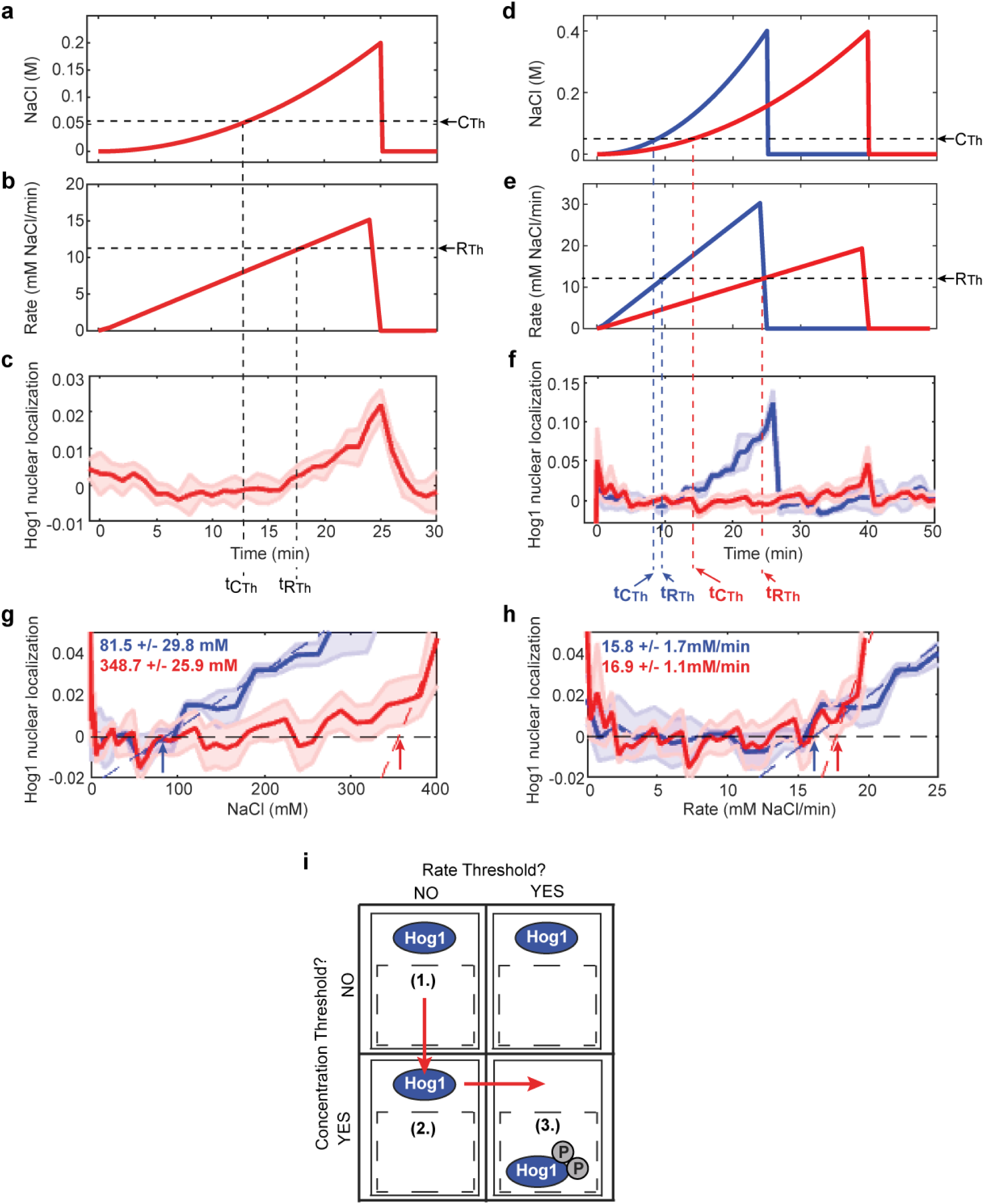
Hog1 nuclear localization timing depends on the rate threshold in quadratic treatments. (**a**) Quadratic NaCl concentration change indicating the time that the concentration threshold (t_CTh_) is satisfied. (**b**) NaCl rate change indicating the time the treatment reaches the rate threshold (t_RTh_). (**c)** Signaling response to the quadratic treatment depicted in “a” and “b” indicating t_CTh_ and t_RTh_. **(d)** Two additional quadratic treatments used to study how treatment rate affects Hog1 translocation. A 25 min quadratic to 0.4 M NaCl is shown in blue and a 40 min quadratic to 0.4 M is shown in red. The concentration threshold (C_Th_) is indicated by the black dotted line. **(e)** Treatment rate during the quadratic treatments depicted in “d”. The rate threshold (R_Th_) is indicated in the black dotted line. **(f)** Hog1 nuclear localization during the quadratic treatments depicted in “d” and “e” plotted as a function of time. The time that each treatment satisfies the C_Th_ and R_Th_ is indicated by the blue dotted lines (for the 25 min quadratic to 0.4 M NaCl) and the red dotted lines (for the 40 min quadratic to 0.4 M NaCl). **(g)** Hog1 activation during the two additional quadratic treatments plotted as a function of concentration. The concentration thresholds correlating with Hog1 nuclear localization during these treatments are indicated in font color coded to match the corresponding treatment. **(h)** Hog1 translocation during the additional quadratic treatments plotted as a function of rate. The rate thresholds correlating with Hog1 activation during these treatments are indicated as in **d**. (**i**) AND logic model for quadratic stimulation.

**Fig. 4.**
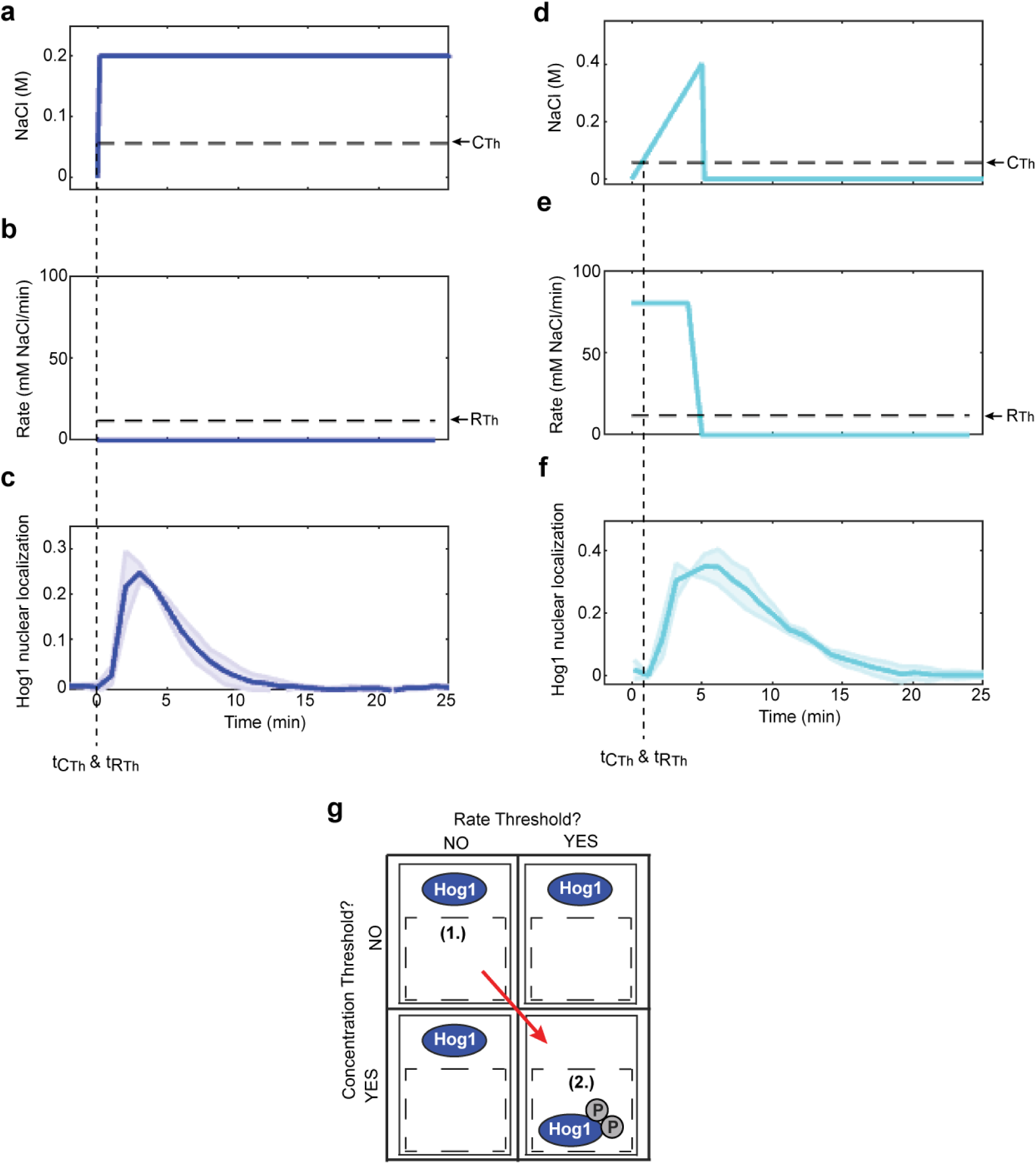
Step and fast ramp stimulations do not enable the detection of threshold conditions. **(a)** Step treatment concentration over time indicating C_Th_. **(b)** Treatment rate during step treatment indicating R_Th_. **(c)** Hog1 nuclear localization during step stimulation indicating the time step treatment satisfied the C_Th_ and R_Th_. **(d)** Fast ramp treatment concentration over time indicating C_Th_. **(e)** Treatment rate during fast ramp treatment indicating R_Th_. **(f)** Hog1 translocation during fast ramp stimulation indicating the time step treatment satisfied the C_Th_ and R_Th_. **(g)** AND logic model indicating the order in which the step and fast ramp treatments satisfy the C_Th_ and R_Th_ conditions.

### Identification of a rate threshold regulator

The existence of a rate threshold raises the possibility of a rate threshold regulator. The ideal regulator would prevent Hog1 from responding when stimulation patterns fail to reach the rate threshold without impinging on the concentration threshold. To identify such a regulator, we first considered key modulators of Hog1 function (Fig. 5a). We specifically focused on the protein tyrosine phosphatases Ptp2 and Ptp3, which act as partially redundant Hog1 off-switches during instantaneous changes in stress concentration^21, 32, 34^, because these factors counteract phosphorylation of the Hog1 tyrosine residue (Tyr-176) required for salt-induced Hog1 activity^44^.

**Fig. 5.**
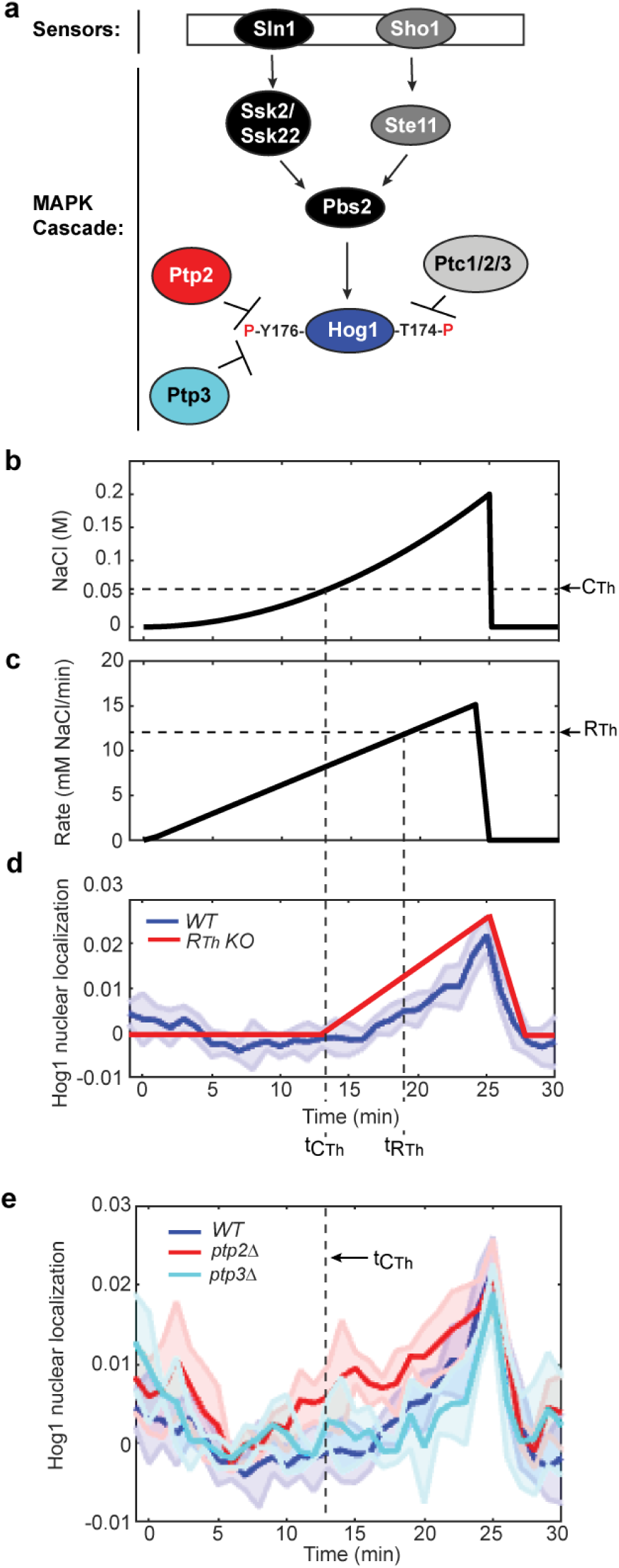
Hog1 localizes to the nucleus at an earlier time point during quadratic treatment in the *ptp2Δ* strain. (**a**) Simplified model of the HOG pathway. Hypothesized regulators of the rate threshold are indicated in red and cyan. (**b**) Quadratic NaCl concentration treatment indicating CTh and t_CTh_. (**c**) NaCl rate during quadratic treatment with RTh and t_RTh_ marked. The treatment rate at the time CTh is satisfied (R at CTh) is also indicated. (**d**) Signaling response to quadratic stimulation in WT (measured, blue) and rate threshold knock out (RTh KO) mutant (predicted, red). t_CTh_ and t_RTh_ are marked. (**e**) Signaling response to quadratic treatment in *WT*, *ptp2*Δ, and *ptp3*Δ cells.

We hypothesized that the deletion of Ptp2 and/or Ptp3 would remove a barrier to stimuli-induced Hog1 activation in stress treatments that exceed the concentration threshold but fall below the rate threshold. To test this hypothesis, we examined Hog1 nuclear accumulation patterns in wild-type (*WT*), *ptp2Δ,* and *ptp3Δ* yeast strains during our quadratic stimulation, in which reduction or removal of the rate threshold would enable translocation earlier in the treatment time course according to the AND logic model (Fig. 5b-d). While the pattern of Hog1 nuclear accumulation in *ptp3Δ* cells appeared similar to *WT* during quadratic stimulation, accumulation in *ptp2Δ* cells occurred much earlier (Fig. 5e).

Reanalyzing accumulation as a function of rate revealed that accumulation in *ptp2Δ* corresponds with a substantially reduced rate (Fig. 6a) compared to *WT*. Further, reanalyzing accumulation as a function of concentration revealed that the timing of nuclear enrichment in *ptp2Δ* corresponds with our measured concentration threshold (Fig. 6b). Thus, in *ptp2Δ* cells, signaling apparently depends only on the concentration threshold condition (Fig. 6c), an effect invisible to pulse treatments in which little difference in translocation pattern was observed among *WT*, *ptp2Δ*, and *ptp3Δ* strains (Fig. 6d-e).

**Fig. 6.**
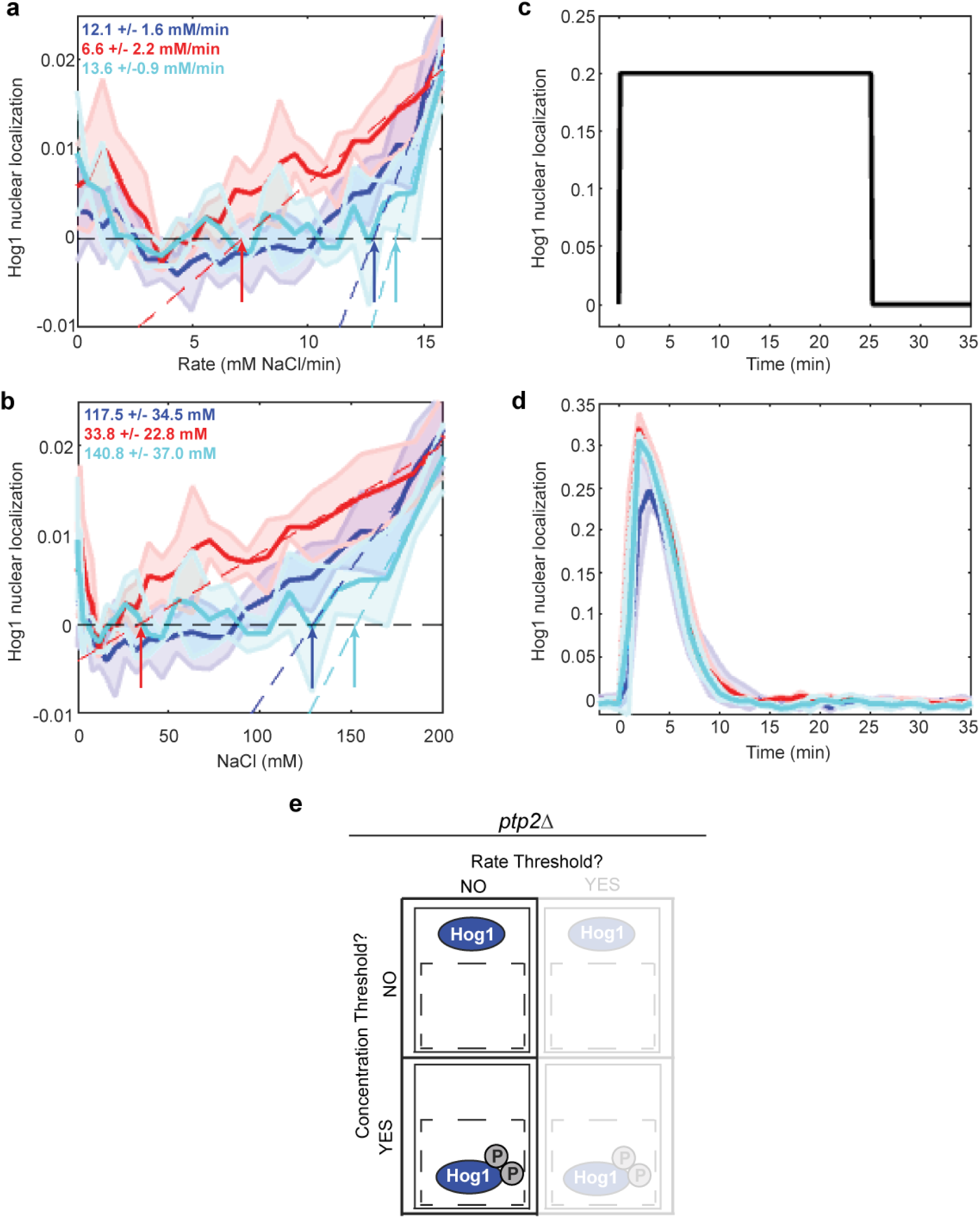
The rate threshold for Hog1 nuclear localization is specifically dependent on the Ptp2 phosphatase. (**a**) Signaling response in the *WT*, *ptp2*Δ, and *ptp3*Δ strains plotted as a function of NaCl treatment rate. The fitted RTh in each strain is listed in bold type. (**b**) Response in each strain plotted as a function of NaCl concentration. The apparent CTh in each strain is listed. **(c)** Step stimulation indicating treatment concentration over time. **(d)** Hog1 translocation in response to the stimulation in “c” in each strain. (**e**) Model of Hog1 activation in the *ptp2*Δ strain indicating the apparent loss of regulation via the rate threshold.

### Tuning the rate threshold

If Ptp2 indeed serves as a major rate threshold regulator, its over-expression should increase the threshold rate required for Hog1 nuclear enrichment. To test this hypothesis, we constitutively expressed *PTP2* in the *ptp2Δ* strain under the control of the upstream regulatory DNA sequences for other genes [*ADH1* and *TEF1*], creating strains with highly increased Ptp2 expression levels (Fig. 7a), and studied Hog1 translocation during quadratic stimulations. A quadratic treatment delivering rates up to 16 mM/min failed to induce activation in the over-expression strains (Fig. 7b-d), while showing activation correspondence with the expected stimulation rates in the empty vector and *PTP2* expression strains (Fig. 7e). However, a quadratic treatment that included rates up to 30 mM/min (Fig. 8a, b) induced Hog1 translocation in all strains (Fig. 8c). Translocation in this second quadratic treatment followed the pattern expected based on the relative timing of the concentration and rate thresholds in the empty vector and native *PTP2* expression strains (Fig. 8d, Supplementary Fig.2), whereas translocation in the over-expression strains corresponded with higher stimulation rates (Fig. 8d). Ptp2 expression levels thus fine-tune the rate required to activate signaling.

**Fig. 7.**
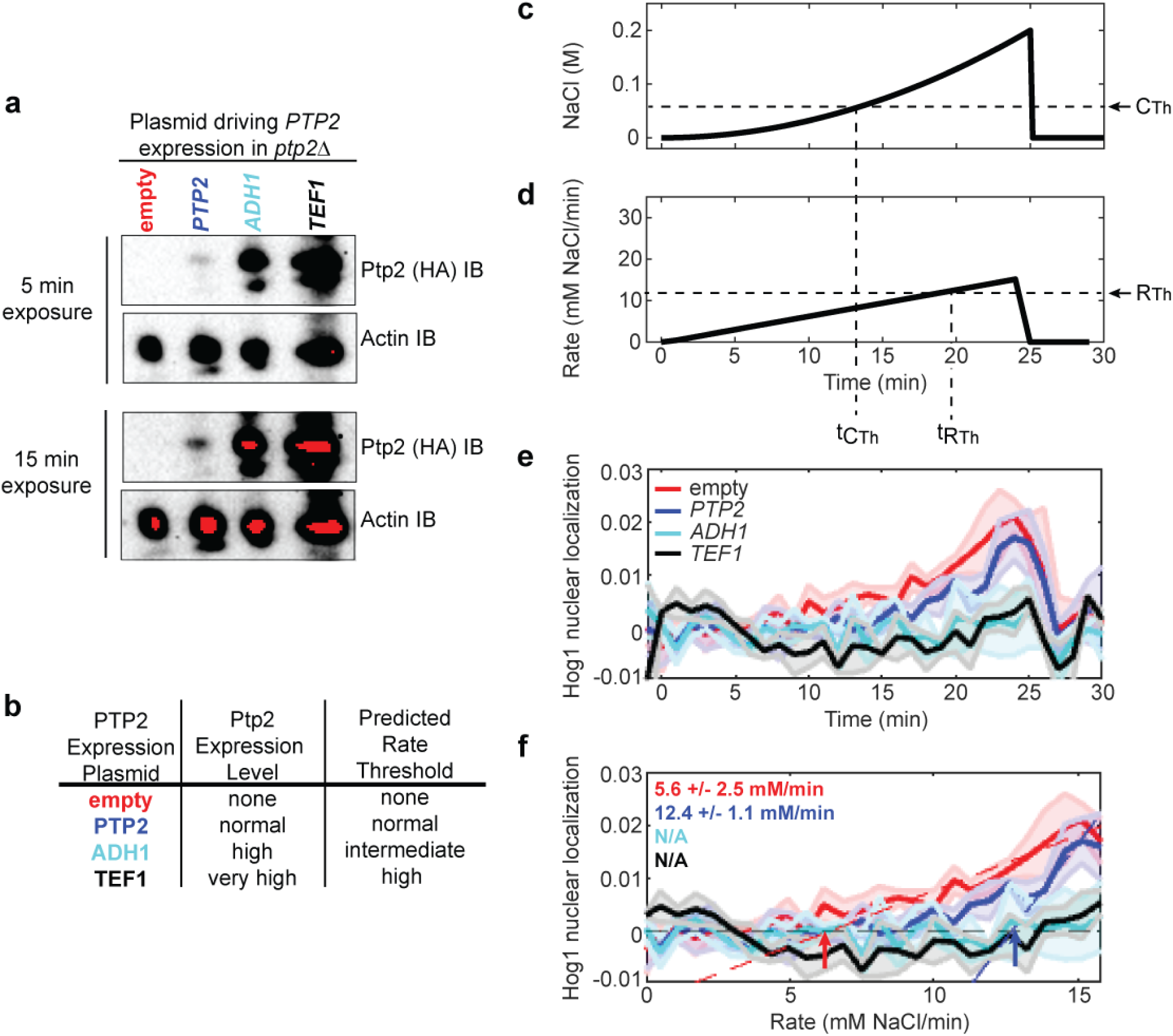
25 minute quadratic treatment to 0.2M NaCl fails to induce Hog1 nuclear localization in the *PTP2* over-expressions strains. (a) HA_3_-tagged Ptp2 expression levels in the *ptp2Δ* strain carrying either an empty vector (empty, red) or one of three plasmids named based on the gene upstream regulatory DNA driving *PTP2* expression [*PTP2* (blue), *ADH1* (cyan), and *TEF1* (black)] were scored via anti-HA immunoblotting relative to loading control (Actin). *Top:* 5 min imaging exposure, *bottom:* 15 min imaging exposure. (**b**) Ptp2 expression levels and predicted rate thresholds in the *ptp2Δ* strain carrying either an empty vector (“empty”) or expressing *PTP2* from the listed expression plasmid. **(c)** Quadratic treatment concentration over time. **(d)** Rate over time for the treatment profile depicted in “b”. **(e)** Hog1 nuclear localization in the *ptp2Δ* strain containing an empty vector (red) or expressing *PTP2* from the upstream regulatory DNA sequences of one of three genes [*PTP2* (blue), *ADH1* (cyan), or *TEF1* (black)]. **(f)** The rate threshold corresponding with Hog1 nuclear localization in each *PTP2* expression variant strain determined using the 25 min quadratic treatment to 0.2M NaCl. The rates corresponding with Hog1 nuclear localization are indicated in bold font in the top left corner of the plot color-coded to the match the corresponding *PTP2* expression variant strain.

**Fig. 8.**
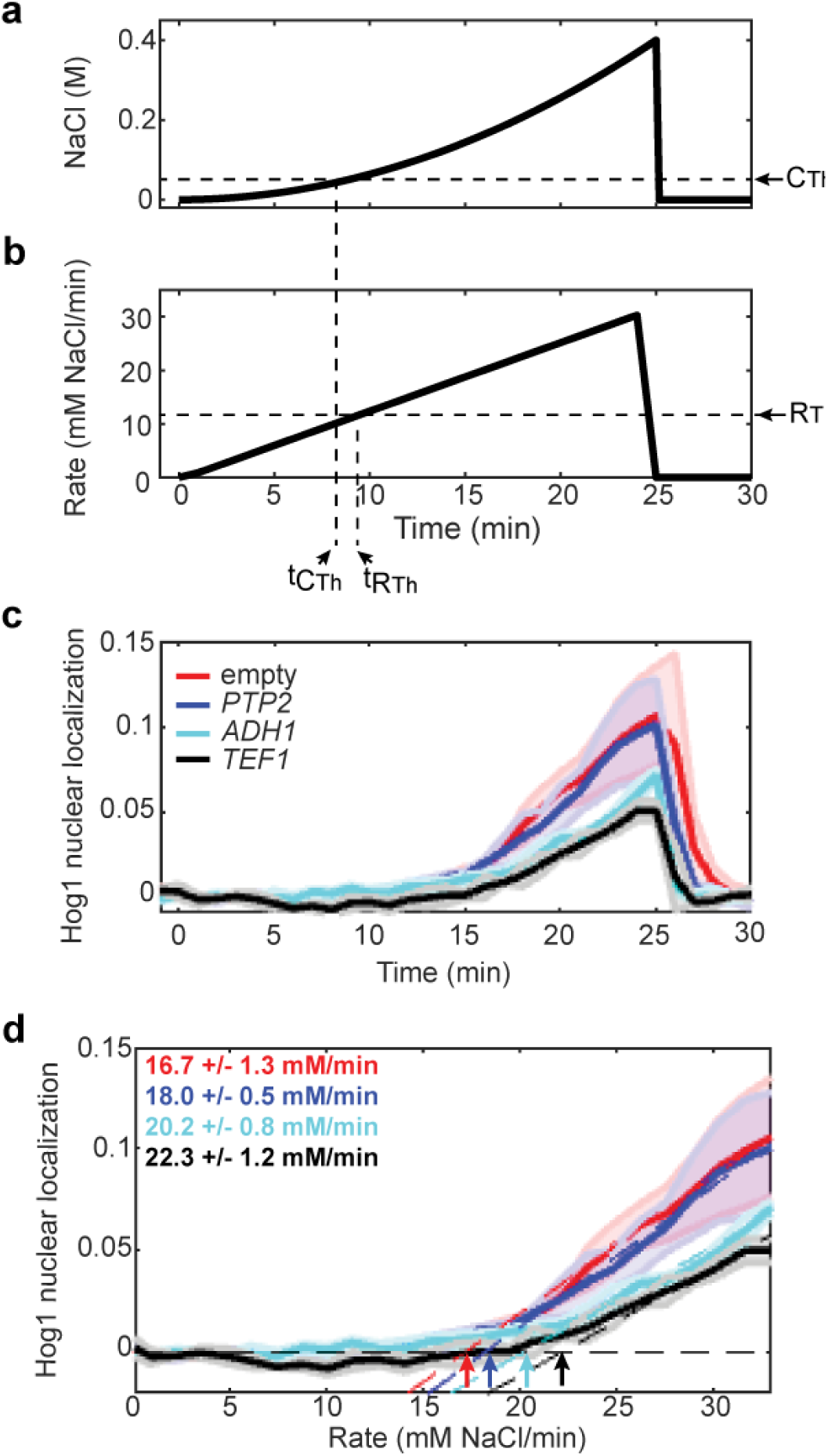
Ptp2 expression levels fine-tune the stimulation rate required to activate Hog1 signaling. (**a**) NaCl concentration during a 0.4M NaCl quadratic stimulation indicating the CTh and t_CTh_. (**b**) NaCl rate during these stimulations indicating RTh and t_RTh_. (**c**) Hog1 nuclear localization response to the quadratic stimulation in “a” and “b” depicted as a function of time in each *PTP2* expression strain. (**d**) Signaling response plotted as a function of NaCl treatment rate. The RTh in each strain are indicated in bold.

In summary, the experiments we have presented directly describe regulation of the HOG pathway by a rate threshold whose set point can be modulated by Ptp2 expression levels. Our rate threshold discoveries represent a paradigm shift in the logic of signaling regulation.

## Discussion

Because stimulation patterns vary in terms of both stimulus concentration and rate of addition in vivo^1–7^, it is important to thoroughly understand how the rate of stimulation affects signal transduction and resulting cell behavior. Here, we have investigated how stimulation rate impacts the budding yeast Hog1 model MAPK pathway. We find that shallow rates of stimulation that fall below a threshold under-prepare cells for future high-stress survival, corresponding with a failure to induce signaling. We also find that this rate threshold collaborates with the concentration threshold in that both conditions must be met for signaling to occur.

These findings demonstrate that the timing of osmostress signaling activation depends as much upon stimulation rate as it does upon stimulus concentration change. Thus, stimulation profile matters at least as much as stimulus identity in determining whether, when, and to what extent signaling and associated cellular activity occur. Using our innovative application of rate-varying stimulus treatments, we find that signal transduction can correspond with either the rate or the concentration threshold depending on the type of stimulus profile applied (Fig. 9). Meanwhile, signaling pathway mutations, such as the *PTP2* deletion and expression level variants, may eliminate or alter the rate threshold requirement.

**Fig. 9.**
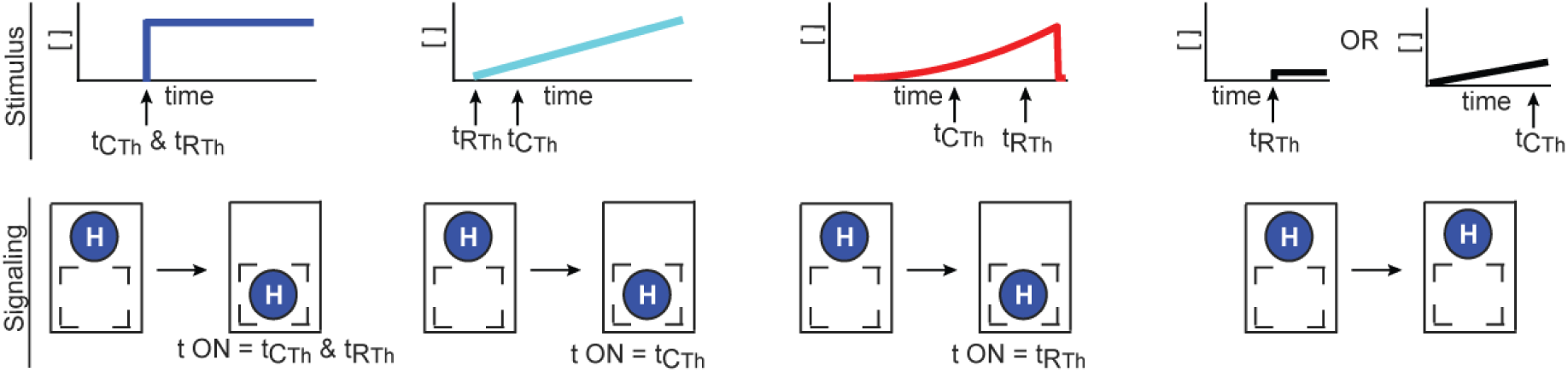
Model of signaling regulation by threshold conditions. The timing of signaling activation (bottom) can correspond with both of the threshold conditions or either the C_Th_ or R_Th_ alone depending on stimulation type (top).

The signaling logic paradigm that controls the yeast osmostress pathway also appears to govern other pathways given the failure of shallow stimulus gradients to trigger *B. subtilus* general stress and mouse myoblast morphogen signaling even when concentrations sufficient for activation during pulsed treatment^10, 12^. We therefore speculate that rate thresholds are a general operating principle of signaling systems, particularly those where known physiological variation in stimulation rate exists^1–7^. A general rate threshold logic mechanism carries with it implications for: (1) how cells make signaling decisions, (2) experimental design, and (3) apparent redundancy in signaling networks.

Rate threshold logic mechanisms likely set important limits on pathway sensitivity to extracellular stimulation. In the case of the HOG pathway studied here, these limits likely serve the same purpose as the sensitivity limits set by the concentration threshold: conserve cellular resources specifically for significant osmostresses. Like pulsed concentration changes below the concentration threshold, slow increases in osmostress may not immediately threaten cell fitness. Consistent with this idea, we do not observe any substantial growth difference between yeast treated with single ramp stimulations above and below the rate threshold (see Supplementary Fig. 1). Rate threshold regulation of osmostress signaling thus appears to ensure an advantageous level of cell sensitivity to stimulation rate. In other pathways, rate threshold mechanisms would provide a helpful way to differentiate signal from noise. The likely rate threshold in the *B. subtilis* general stress pathway^10^ probably also helps conserve resources by activating only when the type of stress the cells experience is unclear. Meanwhile, having cells effectively ignore very shallow morphogen gradients, as appears to be the case in the mouse myoblast morphogen signaling pathway, likely helps direct cells into separate fates during development^11, 12^.

Because both rate and concentration thresholds set the limits of cellular sensitivity to extracellular stimuli, measuring these conditions in other signaling pathways serves as an essential goal in signaling biology. Having these measurements in hand would enable better understanding and prediction of how cells will respond to a given dynamic stimulation profile. Further, defining the limits of cellular sensitivity would enable the detection of changes in these limits caused by signaling pathway mutations such as those common in cancer and developmental disorders^45^. Identifying these aberrant sensitivities would improve our understanding of disease etiology and aid in the design of better treatments. Importantly, the signaling logic paradigm uncovered by our study provides the necessary guide for designing dynamic stimulation profiles that rapidly detect and measure threshold conditions. These profiles are easy to implement and can be readily applied to make threshold condition measurements in any signaling pathway and/or cell type^4, 14^.

The existence of any detected threshold condition begs the question of what regulates the condition. We found that in the yeast HOG pathway, the protein tyrosine phosphatase Ptp2 serves as one major regulator of our newly discovered rate threshold condition. Excitingly, this function provides Ptp2, which is otherwise partially redundant with Ptp3^21, 32, 34^, with an apparently unique role. Additionally, it expands the list of possible roles for phosphatases, which remain much less understood than kinases, in spite of the fact that both types of factors contribute to balancing cellular phosphorylation levels^46^. While phosphatases have generally been thought of as pathway on/off switches, our findings and those of another pioneering study performed using bacterial two-component systems^47^ show that these important signaling factors can regulate cellular sensitivity to extracellular stimulation. Further investigations using dynamic stimulation profiles are needed to determine whether phosphatases in other signaling systems regulate cellular sensitivity to stimulus addition. Such studies may also enable the identification of unique functions for specific phosphatases as well as other factors that appear redundant or partially redundant during switch-like changes in stimulus concentration, thereby facilitating the disentanglement of complex signaling networks.

In conclusion, we have demonstrated a rate threshold mechanism in the HOG pathway along with a highly generalizable strategy for investigating threshold conditions and the signaling pathway components that regulate them. We strongly anticipate that the application of this strategy to other systems will lead to improved disease diagnosis and treatment while transforming our understanding of the logic underlying the complex signaling networks prevalent in higher eukaryotes.

## Methods

### Yeast constructs and strains

All plasmids used in this study (Supplementary Table 1) were constructed via standard restriction enzyme cloning. Appropriate restriction sites were either included in custom synthesized oligonucleotides (IDT) or introduced by PCR with Q5 High-Fidelity Polymerase (NEB) according to the manufacturer’s instructions. All constructs were sequence verified.

The yeast strains used in this study (Supplementary Table 2) were derived from the haploid *Saccharomyces cerevisiae* BY4741 strain^48^. All yeast genomic deletions and integrations were confirmed via PCR. The *PTP2* and *PTP3* genes have been replaced with the kanamycin resistance cassette (*KANMX4*) in the *ptp2*Δ and *ptp3*Δ strains^49^. Strains used to track Hog1-YFP nuclear localization were obtained from a previous study^39^ or generated by C-terminally tagging the endogenous *HOG1* with sequences encoding yellow fluorescent protein (YFP) ^33^ through homologous DNA recombination. Ptp2 expression and empty vector control plasmids were introduced into yeast via high efficiency yeast transformation^50^.

### Yeast growth

Three days before an experiment, yeast cells from a glycerol stock stored at −80 °C were streaked out on Complete Synthetic Media (CSM) (Formedium) or CSM lacking Uracil (CSM-Ura) (Formedium) for selection as needed. The day before the experiment, a single colony from the plate was inoculated in liquid media (pre-culture). After 2 - 6 hr, the optical density at 600 nm (OD_600_) of the pre-culture was measured and diluted into fresh liquid culture to reach an OD_600_ of 0.3-0.5 (depending on the experimental application) the next morning. These OD_600_ values represent 1.5-2.5 x 10^6^ cells/mL.

### Pump profile generation

Pump profiles designed to deliver varying rate salt treatments were generated using an in-house developed MATLAB script that accounts for stock NaCl concentration, pump withdraw rate, volume in the container in which the profile is generated, and culture volume change^14^. Profiles were loaded onto programmable syringe pumps (New Era Pump Systems) prior to treatment.

### High-stress preconditioning assay

High-stress preconditioning assays were designed based on previous work^42, 43^. Initial stress treatments began when cells in culture grown overnight reached an OD_600_ of 0.3. The untreated and 1 hr 0.4 M NaCl pulse 1^st^ stress treatments were chosen as negative and positive controls. Ramp treatments were delivered directly to cells in culture via tubing connecting culture flasks to stock CSM NaCl solution placed in programmable syringe pumps (New Era Pump Systems). These treatments were designed in order to deliver salt at a rate of either 4 mM NaCl/min or 16 mM NaCl/min to a final concentration of 0.4 M NaCl. The time each ramp treatment was held at 0.4 M NaCl was determined such that all first stress treatments that included NaCl received the same cumulative salt exposure (see calculations below). Because treatment duration varied, first stress treatment start times were adjusted such that all treatments ended at the same time. Upon completion of the first treatment, culture OD_600_ was measured and 14mL of each culture was transferred to 15 mL conical tubes, and cells were harvested by centrifugation for 5 min at 3,000 x g. The supernatant was removed, and cells were resuspended in 1 mL sterile H_2_O, transferred to 1.5 mL microcentrifuge tubes (Eppendorf) and pelleted by centrifugation for 5 min at 21,300 x g. Supernatant was removed and pellets were resuspended in 100 µL sterile H_2_O. 5 µL of each resuspended culture was transferred to 145 µL of either CSM (0 M NaCl second stress) or 2 M NaCl CSM (high second stress) in the first well of each row of a 96-well plate. Once all cultures were transferred, each was diluted 1:2 across the wells of each plate column containing 100 µL of each second stress treatment. 96-well plates were incubated at 30 °C with shaking at 250 rpm for 2.5 hrs. After this time, 150 µL CSM was added to each well. Culture in the wells was spotted using a replica plater (Sigma R2508) onto a 15 cm 2% agar CSM plate to enable the determination of relative high-stress survival. To quantitate differences in high second stress survival, culture from the first well of each row (containing the highest concentration of cells from each sample) was diluted 1:2,000, and 600 µL of each treatment was spread onto a 15 cm 2% agar CSM plate.

Plates were incubated for 2-3 days at 30 °C and imaged using a ChemiDoc MP (BioRad). Individual colonies from the plates generated to enable high second stress survival quantitation were counted using in-house developed MATLAB scripts. In brief, after the user is prompted to define the plate diameter by clicking, the scripts process the plate images to generate a uniform background. The image is converted to binary with the background and colonies converted into opposite values and the number of centroids within a user-defined size range in the binary colony value are recorded.

For each first stress treatment, survival was calculated manually by dividing the number of colonies formed from culture exposed to the 2 M NaCl second stress by the number formed from exposure to 0 M NaCl second stress. Relative survival mean and standard deviation were calculated by normalizing the survival resulting from each first stress to mean survival in the untreated first stress culture for three independent biological replicates. Statistical significance was calculated using the Kolmogorov-Smirnov (K-S) test.<u>Example Cumulative Salt Exposure</u>

#### Calculation

-1 hr 0.4 M NaCl step = 60 min X 400 mM NaCl = 24,000 mM NaClmin cumulative salt exposure

-4 mM NaCl/min ramp to 400 mM = (400 mM X 100 min)/2 = 20,0000 mM NaCl min cumulative salt exposure

24,0000 mM NaCl min – 20,000 mM NaClmin = 4,000 mM NaCl min more salt exposure needed

4,000 mMNaCl min/400 mM = 10 min hold after ramp

### Time-lapse microscopy

#### Treatments

3 ml of yeast cells expressing *HOG1-YFP* from the chromosome grown to an OD_600_ of 0.45-0.55 (2.25-2.75 x 10^6^ cells/mL) were pelleted by centrifugation at 500 x g for 6 min. The *HOG1-YFP* strain obtained from a previous study^39^ was used for the experiments presented in Figure 2E, Figure 2F, Figure 3F-H, and Figure 4. The *HOG1-YFP* strains generated for this study were used to collect the data in all other figures. Cell pellets were resuspended in 50 µl CSM and loaded into a flow chamber as previously described^25^.

Three types of perturbations were used in time-lapse microscopy treatments: pulse, linear ramp and quadratic gradients. To deliver the pulse treatments, flow chambers containing cells were connected via tubing to a conical tube containing the CSM NaCl treatment to be delivered on one end and a programmable syringe pump (New Era Pump Systems) set to withdraw media on the other. Ramp and quadratic perturbations were set up as described previously^14^. In brief, these perturbations were generated in a beaker containing CSM subjected to constant mixing on a stir plate by delivering CSM NaCl stock solution in pre-programmed steps via syringe pump (New Era Pump Systems). Media was drawn from this beaker to the flow chamber over the time course of treatment profile generation using a syringe pump programmed to withdraw media at a rate of 0.1 mL/min. At the completion of all NaCl treatments, cells were returned to CSM by switching the beaker or conical containing CSM NaCl to a conical containing CSM only.

#### Image acquisition

The Micro-Manager program (https://micro-manager.org/) version 1.4 was used to control a Nikon T*i* Eclipse epi-fluorescent microscope equipped with perfect focus (Nikon), a 100x VC DIC lens (Nikon), a fluorescent filter for YFP (Semrock), an X-cite fluorescent light source (Excelitas) and an Orca Flash 4v2 CMOS camera (Hamamatsu). YFP images were collected every 1 min with an exposure of 20 ms. Bright-field (TRANS) images with an exposure of 10 ms were collected every 10 sec during each time-lapse experiment. 3-4 YFP images of cells loaded on the flow chamber were taken prior to the start of each treatment to enable Hog1 nuclear localization baseline level determination.

#### Image analysis

Image segmentation was performed in two-steps. First, YFP images (detecting Hog1-YFP) were smoothed and background corrected. A threshold was set to identify the brightest pixels for each cell. The region containing the 100 brightest pixels in each cell was then used as an intracellular marker. Second, TRANS images were smoothed, background corrected and overlain with the corresponding YFP image (or the YFP image from the previous time point in the case were no YFP image was taken). A watershed algorithm was applied to these overlain images to segment and label the objects (cells). After segmentation, objects that deviate significantly in size from the average cell or on the border of the image are removed, resulting in an image with segmented cells. This process was repeated for each image. After segmentation, the centroid of each cell was computed and stored. The distance between the centroids of each cell in consecutive images were compared. The centroids that show the smallest distance difference between two consecutive images are considered the same cell at two different time points. This whole procedure is repeated for each image resulting in single cell trajectories. Each single cell trace was visually inspected, and cells exhibiting large fluctuations in YFP or TRANS signal over the time course were removed. In a typical experiment, 20-30% of a starting population of 200-450 single cells remained after this filtering step for further processing.

To determine Hog1 nuclear localization in each cell, the average per pixel fluorescent intensity of the whole cell (I_w_) and of the top 100 brightest fluorescent pixels (I_t_) was calculated. In addition, fluorescent signal per pixel of the camera background (I_b_) was recorded. Hog1 nuclear localization was calculated as Hog1(t) = [(I_t_(t) - I_b_) / (I_w_(t) - I_b_)]. The value for Hog1(t = 0) defines Hog1 nuclear localization baseline level. Single cell traces were smoothed and nuclear localization over the time course was determined by subtracting Hog1(t=0) from Hog1(t). For cell volume measurements, volume change relative to the volume at the beginning of the experiment was calculated. For both the single cell volume and Hog1(t) fluorescent traces, the median and the average median distance (the equivalent of the standard deviation if the median is used instead of the mean) were computed to put less weight on sporadic outlier cells due to the image segmentation process.

The in-house developed codes used for image segmentation^51^ and time-lapse image analysis^14, 40, 52^ have been previously described.

### Determination of concentration and rate thresholds

The rate and concentration thresholds were determined by plotting the median Hog1 nuclear localization in each biological replicate as a function of treatment concentration or rate, respectively. Two points were manually selected from each of these plots demarcating the beginning and end point of robust Hog1 nuclear enrichment. The data between these points was automatically fit to the linear function (y = mx + b). The “x” value (concentration or rate) at the intersection point of a horizontal line at Hog1 = 0 (baseline) and the line fitting Hog1 nuclear enrichment pattern (activation) was determined. The average and standard deviation of the “x” values from the replicates for each treatment profile represent the concentration and rate thresholds corresponding with Hog1 nuclear localization in those treatment profiles. The average slope (“m”) and y-intercept (“b”) from the linear function fit to the Hog1 activation data for each biological replicate obtained from a particular treatment profile were used to generate the dotted lines overlaying Hog1 nuclear enrichment in the threshold determination figures.

### Immunoblotting

Mild alkali treatment and heating in standard electrophoresis buffer^53^ were used to extract total protein from yeast cell pellets containing 1 OD_600_ unit (5 x 10^6^ cells). Proteins were fractionated on a NuPAGE 4-12% Bis-Tris denaturing polyacrylamide gel (Thermo Fisher Scientific). Gels were equilibrated in transfer buffer (30 mM Bicine, 25 mM Bis-Tris, 1 mM EDTA, 60 µM chlorobutanol) and electrotransferred to PVDF membranes pre-equilibrated in transfer buffer (GE, Immobilon-P. 0.45µM). 5% w/v non-fat milk in 1 X Tris-buffered saline (TBS) (100mM Tris-Cl (pH 7.5), 150 mM NaCl) was used for blocking. HAx3-Ptp2 was detected using an anti-HA HRP conjugate 3F10 (Roche) at a 1:5,000 dilution. Endogenous actin (used as a loading control) was detected using a 1:5,000 dilution of anti-ß-actin antibody (Abcam #ab8224) followed by incubation with a 1:2,500 dilution of HRP-conjugated horse anti-mouse IgG antibody (Cell Signaling #7076S). All antibody incubations were performed in 1 % w/v non-fat milk blocking in 1X TBS. Antibody signals were visualized using. Enhanced chemiluminescence (ECL) reagent (GE) was used to detect antibody signals. Signals were visualized using a ChemiDoc MP Imaging System (BioRad).

### Data availability

The datasets generated during and/or analysed during the current study are available from the corresponding author on reasonable request.

### Code availability

Cell segmentation codes have been previously described^51^ and are available in the public dropbox repository: https://www.dropbox.com/sh/egb27tsgk6fpixf/AADaJ8DSjab_c0gU7N7ZF0Zba?dl=0.

Image processing codes have also been previously described^14, 40, 52^. All codes used for image processing and analysis in this study will be made available upon reasonable request.

## Acknowledgements

We would like to thank Mr. Cleland, Mr. Hughes, Mr. Venkat and Drs. Weil, Tansey, Wadzinski, Carrasco, and Colbran for feedback on the study and manuscript. We would also like to thank the Vanderbilt Molecular Biology core facility for providing access to and technical assistance with instrumentation necessary to complete this work. This work was supported by the NIH Director’s New Innovator Award (1DP2GM114849), the NIGMS (5R01GM115892), the Advanced Computing Center for Research and Education (ACCRE) (1S10OD023680-01), and Vanderbilt University Startup Funds. AJ was supported by the Ion Channels and Transporters Training Grant (T32NS007491), BK was supported by the Molecular Biophysics Training Grant (5T32GM008320) and AT was supported by an AHA predoctoral fellowship (18PRE34050016). The content of this publication is solely the responsibility of the authors and does not necessarily represent the official views of the National Institutes of Health.

## Author contributions

Conceptualization: A.N.J. and G.N. Data Curation: A.N.J., H.J., and G.N., Formal Analysis: A.N.J., H.J., and G.N., Funding Acquisition: G.N. Investigation: A.N.J., G.L., H.J., A.T., Methodology: A.N.J., G.L., H.J., and A.T., Project administration: A.N.J. and G.N., Resources: A.N.J., D.R., and G.N., Software: B.K., H.J., and G.N., Supervision: G.N., Validation: A.N.J. and H.J., Visualization: A.N.J., Writing-original draft: A.N.J., Writing-reviewing and editing: all authors.

## Additional Information

**Supplementary Information** is available for this paper

**Correspondence and requests for materials** should be addressed to G.N.

## Main References

1. Polonsky, K. S., Given, B. D. & Van Cauter, E. Twenty-four-hour profiles and pulsatile patterns of insulin secretion in normal and obese subjects. J. Clin. Invest. 81, 442–448 (1988).

2. Hoffmann, E. K., Lambert, I. H. & Pedersen, S. F. Physiology of Cell Volume Regulation in Vertebrates. Physiol. Rev. 89, 193–277 (2009).

3. Chovatiya, R. & Medzhitov, R. Stress, Inflammation, and Defense of Homeostasis. Mol. Cell 54, 281–288 (2014).

4. Shimizu, T. S., Tu, Y. & Berg, H. C. A modular gradient-sensing network for chemotaxis in Escherichia coli revealed by responses to time-varying stimuli. Mol. Syst. Biol. 6, 382 (2010).

5. Wang, C. J., Bergmann, A., Lin, B., Kim, K. & Levchenko, A. Diverse Sensitivity Thresholds in Dynamic Signaling Responses by Social Amoebae. Sci. Signal. 5, 213 (2012).

6. Brabant, G., Prank, K. & Schofl, C. Pulsatile patterns in hormone secretion. Trends Endocrinol. Metab. 3, 183–190 (1992).

7. Sagner, A. & Briscoe, J. Morphogen interpretation: concentration, time, competence, and signaling dynamics. Wiley Interdiscip. Rev. Dev. Biol. 6, e271 (2017). doi:10.1002/wdev.271

8. Forsten, K. E. & Lauffenburger, D. A. Autocrine ligand binding to cell receptors. Mathematical analysis of competition by solution ‘decoys’. Biophys. J. 61, 518–529 (1992).

9. Ang, J., Harris, E., Hussey, B. J., Kil, R. & McMillen, D. R. Tuning response curves for synthetic biology. ACS Synth. Biol. 2, 547–567 (2013).

10. Young, J. W., Locke, J. C. W. & Elowitz, M. B. Rate of environmental change determines stress response specificity. Proc. Natl. Acad. Sci. 110, 4140–4145 (2013).

11. Heemskerk, I. et al. Rapid changes in morphogen concentration control self-organized patterning in human embryonic stem cells. Elife 8, e40526 (2019).

12. Sorre, B., Warmflash, A., Brivanlou, A. H. H. & Siggia, E. E. D. Encoding of temporal signals by the TGF-β Pathway and implications for embryonic patterning. Dev. Cell 30, 334–342 (2014).

13. Kubota, H. et al Temporal Coding of Insulin Action through Multiplexing of the AKT Pathway. Mol. Cell 46, 820–832 (2012).

14. Thiemicke, A., Jashnsaz, H., Li, G. & Neuert, G. Generating kinetic environments to study dynamic cellular processes in single cells. Sci. Rep. 9, 10129 (2019).

15. Fujita, K. A et al. Decoupling of Receptor and Downstream Signals in the Akt Pathway by Its Low-Pass Filter Characteristics. Sci. Signal. 3, ra56 (2010).

16. Sasagawa, S., Ozaki, Y., Fujita, K. & Kuroda, S. Prediction and validation of the distinct dynamics of transient and sustained ERK activation. Nat. Cell Biol. 7, 365–373 (2005).

17. Goulev, Y. et al. Nonlinear feedback drives homeostatic plasticity in H _2_ O _2_ stress response. Elife 6, e2397 (2017).

18. Brewster, J. L. & Gustin, M. C. Hog1: 20 years of discovery and impact. Sci. Signal. 7, re7 (2014).

19. Brewster, J., de Valoir, T., Dwyer, N., Winter, E. & Gustin, M. An osmosensing signal transduction pathway in yeast. Science 259, 1760–1763 (1993).

20. Westfall, P. J. When the Stress of Your Environment Makes You Go HOG Wild. Science 306, 1511–1512 (2004).

21. Saito, H. & Posas, F. Response to Hyperosmotic Stress. Genetics 192, 289–318 (2012).

22. Ferrigno, P., Posas, F., Koepp, D., Saito, H. & Silver, P. A. Regulated nucleo/cytoplasmic exchange of HOG1 MAPK requires the importin beta homologs NMD5 and XPO1. EMBO J. 17, 5606–14 (1998).

23. English, J. G. et al. MAPK feedback encodes a switch and timer for tunable stress adaptation in yeast. Sci. Signal. 8, ra5 (2015).

24. Reiser, V., Ruis, H. & Ammerer, G. Kinase activity-dependent nuclear export opposes stress-induced nuclear accumulation and retention of Hog1 mitogen-activated protein kinase in the budding yeast Saccharomyces cerevisiae. Mol. Biol. Cell 10, 1147–1161 (1999).

25. Muzzey, D., Gómez-Uribe, C. A., Mettetal, J. T. & van Oudenaarden, A. A Systems-Level Analysis of Perfect Adaptation in Yeast Osmoregulation. Cell 138, 160–171 (2009).

26. Macia, J. et al. Dynamic Signaling in the Hog1 MAPK Pathway Relies on High Basal Signal Transduction. Sci. Signal. 2, ra13 (2009).

27. Granados, A. A. et al. Distributing tasks via multiple input pathways increase cellular survival in stress. Elife 6, e21415 (2017).

28. Mitchell, A., Wei, P. & Lim, W. A. Oscillatory stress stimulation uncovers an Achilles heel of the yeast MAPK signaling network. Science 350, 1379–1383 (2015).

29. Hersen, P., McClean, M. N., Mahadevan, L. & Ramanathan, S. Signal processing by the HOG MAP kinase pathway. Proc. Natl. Acad. Sci. U. S. A. 105, 7165–70 (2008).

30. Westfall, P. J., Patterson, J. C., Chen, R. E. & Thorner, J. Stress resistance and signal fidelity independent of nuclear MAPK function. Proc. Natl. Acad. Sci. U. S. A. 105, 12212–12217 (2008).

31. Westfall, P. J. & Thorner, J. Analysis of Mitogen-Activated Protein Kinase Signaling Specificity in Response to Hyperosmotic Stress: Use of an Analog-Sensitive HOG1 Allele. Eukaryot. Cell 5, 1215–1228 (2006).

32. Wurgler-Murphy, S. M., Maeda, T., Witten, E. A. & Saito, H. Regulation of the Saccharomyces cerevisiae HOG1 mitogen-activated protein kinase by the PTP2 and PTP3 protein tyrosine phosphatases. Mol. Cell. Biol. 17, 1289–97 (1997).

33. Warmka, J., Hanneman, J., Lee, J., Amin, D. & Ota, I. Ptc1, a Type 2C Ser/Thr Phosphatase, Inactivates the HOG Pathway by Dephosphorylating the Mitogen-Activated Protein Kinase Hog1. Mol. Cell. Biol. 21, 51–60 (2001).

34. Mattison, C. P. & Ota, I. M. Two protein tyrosine phosphatases, Ptp2 and Ptp3, modulate the subcellular localization of the Hog1 MAP kinase in yeast. Genes Dev. 14, 1229–1235 (2000).

35. Maeda, T., Wurgler-Murphy, S. M. & Saito, H. A two-component system that regulates an osmosensing MAP kinase cascade in yeast. Nature 369, 242–245 (1994).

36. Young, C., Mapes, J., Hanneman, J., Al-Zarban, S. & Ota, I. Role of Ptc2 type 2C Ser/Thr phosphatase in yeast high-osmolarity glycerol pathway inactivation. Eukaryot. Cell 1, 1032–40 (2002).

37. Jacoby, T. et al. Two protein-tyrosine phosphatases inactivate the osmotic stress response pathway in yeast by targeting the mitogen-activated protein kinase, Hog1. J. Biol. Chem. 272, 17749–17755 (1997).

38. Martín, H., Flández, M., Nombela, C. & Molina, M. Protein phosphatases in MAPK signalling: we keep learning from yeast. Mol. Microbiol. 58, 6–16 (2005).

39. Mettetal, J. T. et al. The Frequency Dependence of Osmo-Adaptation in Saccharomyces cerevisiae. Science 319, 482–484 (2008).

40. Neuert, G. et al. Systematic Identification of Signal-Activated Stochastic Gene Regulation. Science 339, 584–587 (2013).

41. Lewis, J. G., Learmonth, R. P. & Watson, K. Induction of heat, freezing and salt tolerance by heat and salt shock in Saccharomyces cerevisiae. Microbiology 141, 687–694 (1995).

42. Berry, D. B. & Gasch, A. P. Stress-activated Genomic Expression Changes Serve a Preparative Role for Impending Stress in Yeast. Mol. Biol. Cell 19, 4580–4587 (2008).

43. Berry, D. B. et al. Multiple Means to the Same End: The Genetic Basis of Acquired Stress Resistance in Yeast. PLoS Genet. 7, e1002353 (2011).

44. Bell, M. & Engelberg, D. Phosphorylation of Tyr-176 of the Yeast MAPK Hog1/p38 Is Not Vital for Hog1 Biological Activity. J. Biol. Chem. 278, 14603–14606 (2003).

45. Lim, W., Mayer, B. & Pawson, T. Cell signaling : principles and mechanisms. (Garland Science, 2015).

46. Fahs, S., Lujan, P. & Köhn, M. Approaches to Study Phosphatases. ACS Chem. Biol. 11, 2944–2961 (2016).

47. Landry, B. P., Palanki, R., Dyulgyarov, N., Hartsough, L. A. & Tabor, J. J. Phosphatase activity tunes two-component system sensor detection threshold. Nat. Commun. 9, 1433 (2018).

48. Brachmann, C. B. et al. Designer deletion strains derived from Saccharomyces cerevisiae S288C: A useful set of strains and plasmids for PCR-mediated gene disruption and other applications Yeast 14, 115–132 (1998).

49. Winzeler, E. A. et al. Functional characterization of the S. cerevisiae genome by gene deletion and parallel analysis. Science 285, 901–906 (1999).

50. Gietz, R. D. & Woods, R. A. Transformation of yeast by lithium acetate/single-stranded carrier DNA/polyethylene glycol method. Methods Enzymol. 350, 87–96 (2002).

51. Kesler, B., Li, G., Thiemicke, A., Venkat, R. & Neuert, G. Automated cell boundary and 3D nuclear segmentation of cells in suspension. Sci. Rep. 9, 10237 (2019).

52. Munsky, B., Li, G., Fox, Z. R., Shepherd, D. P. & Neuert, G. Distribution shapes govern the discovery of predictive models for gene regulation. Proc. Natl. Acad. Sci. 115, 7533–7538 (2018).

53. Kushnirov, V. V. Rapid and reliable protein extraction from yeast. Yeast 16, 857–860 (2000).

