## Supplementary_Information for "A rate threshold mechanism regulates MAPK stress signaling and survival"

**This PDF file includes:**

Figs. S1 to S2
Tables S1 to S2
Supplementary References

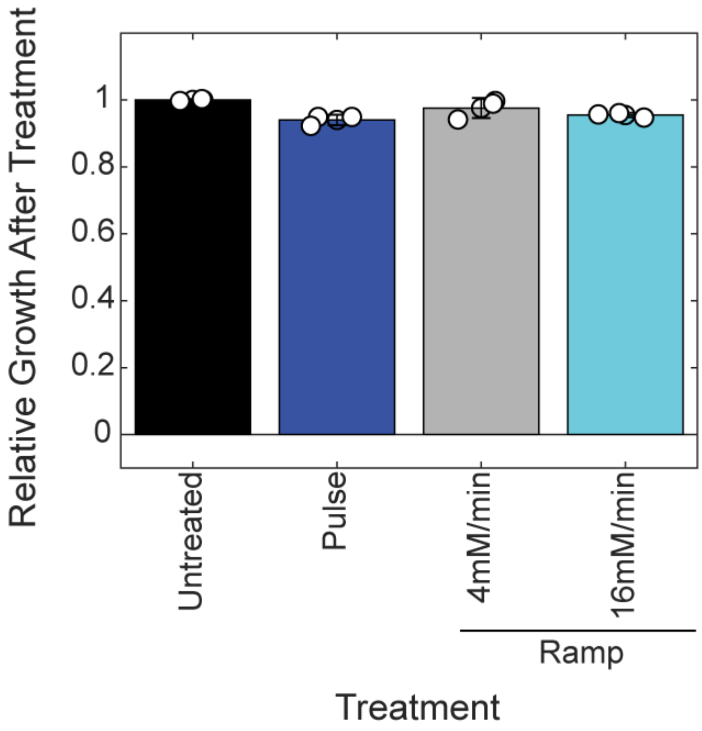


**Supplementary Figure 1. Treatments delivered at rates below and above the rate threshold show similar growth levels after treatment delivery.** Growth relative to untreated culture as determined by measuring the $OD_{600}$ of cultures started at the same $OD_{600}$ after treatment. Pulse and 16 mM/min ramp treatments exceed the rate threshold, while the 4 mM/min ramp treatment does not meet the rate threshold condition. Bars and errors are means and standard deviations of three biological replicates.

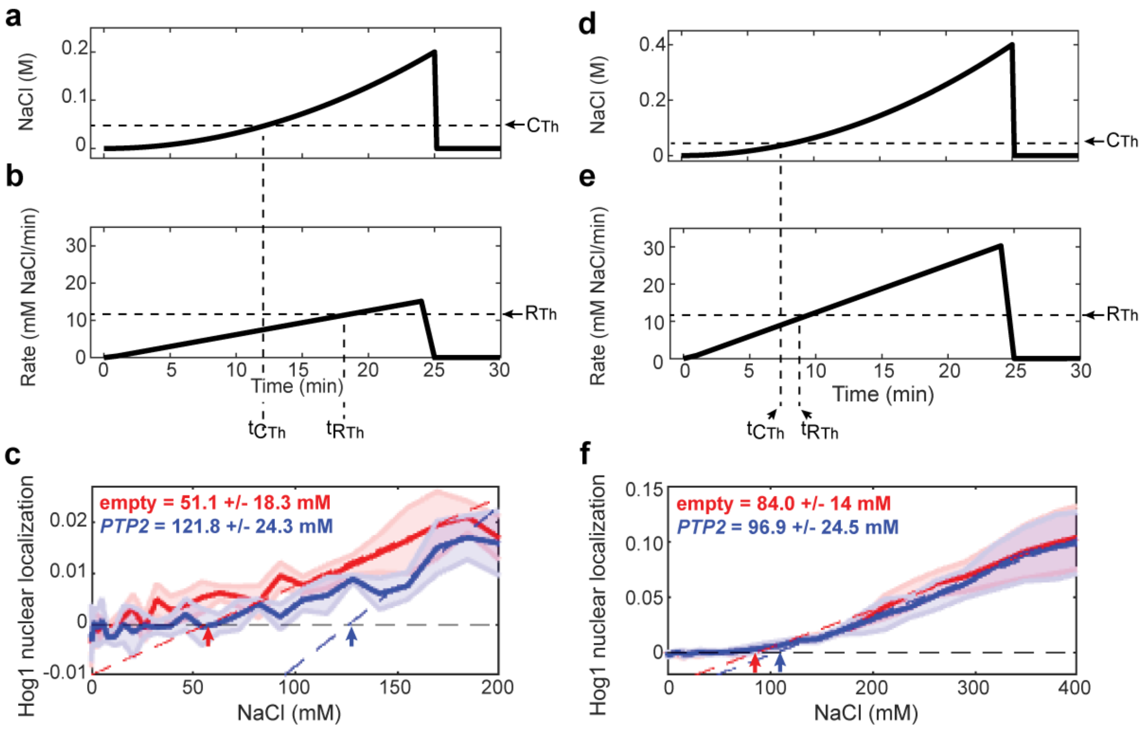


**Supplementary Figure 2. The time between threshold conditions impacts the difference between apparent concentration thresholds. (a)** 25 min quadratic treatment to 0.2M NaCl showing the $C_{Th}$. **(b)** Treatment rate during the quadratic treatment shown in **a**. **(c)** Hog1 activation during the quadratic treatment shown in **a** and **b** plotted as a f-unction of treatment concentration in the *ptp2Δ* strain containing either an empty vector (red) or a plasmid driving *PTP2* using its native upstream regulatory DNA sequences (*PTP2*, blue). The concentration thresholds correlating with Hog1 nuclear localization in each strain during this treatment are indicated in the upper left-hand corner of the plot. **(d)** 25 min quadratic treatment to 0.4 M NaCl showing the $C_{Th}$. **(e)** Treatment rate during the quadratic treatment shown in **d**. **(f)** Hog1 nuclear localization during the quadratic treatment shown in **d** and **e** plotted as a function of treatment concentration as in **c**.

**Supplementary Table 1 – Plasmids used in this study**

| Name | Purpose | Reference |
|---|---|---|
| pRS416 | Empty vector control | 1 |
| pRS416 $UAS_{Ptp2}$ $HA_3$ *PTP2* | WT *PTP2* expression | This study |
| pRS416 $UAS_{Adh1}$ $HA_3$ *PTP2* | Increased *PTP2* expression | This study |
| pRS416 $UAS_{Tef1}$ $HA_3$ *PTP2* | Increased *PTP2* expression | This study |

**Supplementary Table 2 – Yeast strains used in this study**

| Strain | Genotype | Reference |
|---|---|---|
| BY4741 | *MAT***a** *his3Δ1 leu2Δ0 met15Δ0 ura3Δ0* | 2 |
| Hog1-YFP #1 | *MAT***a** *his3Δ1 leu2Δ0 met15Δ0 ura3Δ0 HOG1-YFP::HIS3 NRD1-mRFP1.3* $UAS_{MYO2}$*-rtTA* | 3 |
| Hog1-YFP #2 | *MAT***a** *his3Δ1 leu2Δ0 met15Δ0 ura3Δ0 HOG1-YFP::HIS3* | This study |
| Ptp2Δ Hog1-YFP | *MAT***a** *his3Δ1 leu2Δ0 met15Δ0 ura3Δ0 ptp2Δ::KANMX4 HOG1-YFP::HIS3* | This study |
| Ptp3Δ Hog1-YFP | *MAT***a** *his3Δ1 leu2Δ0 met15Δ0 ura3Δ0 ptp3Δ::KANMX4 HOG1-YFP::HIS3* | This study |
| Ptp2Δ Hog1-YFP + empty | *MAT***a** *his3Δ1 leu2Δ0 met15Δ0 ura3Δ0 ptp2Δ::KANMX4* (pRS416) *HOG1-YFP::HIS3* | This study |
| Ptp2Δ Hog1-YFP + Ptp2 | *MAT***a** *his3Δ1 leu2Δ0 met15Δ0 ura3Δ0 ptp2Δ::KANMX4* (pRS416 $UAS_{Ptp2}$ $HA_3$ *PTP2*) *HOG1-YFP::HIS3* | This study |
| Ptp2Δ Hog1-YFP + Adh1 | *MAT***a** *his3Δ1 leu2Δ0 met15Δ0 ura3Δ0 ptp2Δ::KANMX4* (pRS416 $UAS_{Adh1}$ $HA_3$ *PTP2*) *HOG1-YFP::HIS3* | This study |
| Ptp2Δ Hog1-YFP + Tef1 | *MAT***a** *his3Δ1 leu2Δ0 met15Δ0 ura3Δ0 ptp2Δ::KANMX4* (pRS416 $UAS_{Tef1}$ $HA_3$ *PTP2*) *HOG1-YFP::HIS3* | This study |

**Supplementary References**


1. D. Mumberg, R. Müller, M. Funk, Yeast vectors for the controlled expression of heterologous proteins in different genetic backgrounds. *Gene*. **156**, 119–122 (1995).
2. C. B. Brachmann *et al.*, Designer Deletion Strains derived from Saccharomyces cerevisiae S288C: a Useful set of Strains and Plasmids for PCR-mediated Gene Distruption and Other Applications. *Yeast*. **14**, 115–132 (1998).
3. J. T. Mettetal *et al.*, The Frequency Dependence of Osmo-Adaptation in Saccharomyces cerevisiae. *Science (80-. )*. **319**, 482–484 (2008).
